# Scratching bouts are modeled as Bernoulli trials until successful itch-extinguishing in mice

**DOI:** 10.1101/2022.02.03.478919

**Authors:** Kotaro Honda, Mitsutoshi Tominaga, Kenji Takamori

## Abstract

Itching and subsequent scratching behavior have been observed in many species, including humans. The behavior was evolved to remove skin parasites. Yet, scratching is performed without reliable indicators of whether a parasite is present. We addressed this apparent paradox by studying scratching in mice. Video recordings of ~5000 scratching bouts were collected in free-moving C57BL6/J mice. The statistical properties of their temporal sequence were analyzed. Inter-bout time intervals preceding over 50% of 5000 bouts were <10 s. We hypothesized that episodes of repetitive scratching corresponded to the duration of discrete events of itch sensation and comprised bouts separated by inter-bout intervals of <10 s. The distribution of itch episodes comprising n (n = 1, 2, 3, …) scratching bouts was well-approximated by the geometric distribution with success probability = 0.5 in healthy mice and lower probability in dry skin mice. This suggests that scratching bouts are modeled by probabilistic Bernoulli trials, and their repetitive sequence in each episode continues until the itch is successfully extinguished. Accordingly, we can presume the presence of parasites from the repeat length of scratching bouts determined by the probability of successful itch-extinguishing. This may provide a promising stochastic model to assess itchy phenotypes.

## Introduction

Itch is an unpleasant sensation that triggers an urge to scratch the skin in the affected area and has been documented in many animal species, including turtles (Mui et al., 2012). The itch sensation may have evolved to inform the animal of a potential danger posed by skin parasites, chemical irritants, etc. For example, when we feel itchy in our own back, we scratch there and ask our housemates whether the area scratched is reddened. Physiological scratching bouts in healthy state is explained as a kind of grooming behaviors that maintain the skin homeostasis (Sachs, 1988), and pathological scratching bouts often destroy the corneum barrier of the skin and delay the recovery from skin diseases. Itch is common symptoms in wide variety of diseases such as allergic skin diseases, visceral diseases, cancer, anxiety disorder (Yosipovitch & Bernhard, 2013), while the qualitative difference of itch between these symptoms and healthy state is unclear. Because the patients are suffering form not only damaging the skin but also sleep disorder (Ramirez et al., 2019) and loss of motivation to study and work by itch (Misery et al., 2018), which result in significant economic burden (Girolomoni et al., 2021), we need to control pathological itch emotion. However, it was difficult to objectively understand how unbearable the itch was, and a method for quantifying the quality of the emotion of itch has not been developed because of the complexity of scratching (Wimalasena et al., 2021).

The physiological significance of scratching bouts is still unknown, but a possible theory is that it plays a role in removing skin parasites such as mites (Hart, 1990). However, skin parasites are smaller than the size that can be recognized by the tactile and visual senses. It is actually impossible to know whether parasites exist on the itchy skin, and it is also impossible to confirm whether parasites have disappeared after the scratching bouts, so there seems to be no point in repeating scratching bouts. The emotion of itch, therefore, has never been studied from the perspective of a question “why do we repeat scratching bouts when we are itchy?”

Independent trials with two possible outcomes (such as success or failure) at constant probability are called Bernoulli trials. When the Bernoulli trials are repeated until getting one success, the times of repeated trials describe characteristic probability distribution, which is called the geometric distribution. In the theory of achievement motivation, the subjective success probability affects our motivation to achieve success, especially the probability 0.5 brings out the best motivation in human (Atkinson, 1957). However, it has never been reported how these probability distribution and subjective success probability contribute to maintaining homeostasis and the impulsive emotional behaviors, such as scratching.

We hypothesized that the pattern of the number of repeats of scratching bouts for duration of itch emotion represents the quality of itchiness. To analyze the pattern of the repeats, we determined the first and repeated scratching bouts for duration of the emotion of itch, using approximately 5,000 scratching bouts data obtained from 24-hour recording of healthy mice with back 2 × 2 cm^2^ shaved, under physiological conditions. As a result, we found that the proportion of itch episodes in which the number of repeats of scratching bouts was 1, 2, 3, … was close to (0.5)^1^, (0.5)^2^, (0.5)^3^, … both in male and female mice. This was comparable to a graph of the geometric distribution with the success probability at 0.5, and indicated that repeats of the scratching bouts were the Bernoulli trials until the emotion of itch was extinguished. There has never been a report on the principle that animals use probability to determine the number of repetitive actions, but some models exist (Wang, 2002). We then devised a stochastic control model, in which two outcomes can be obtained stochastically by simultaneous excitation of mutual inhibitory neurons in the central nervous system (Kogo et al., 2021).

Results of the present study suggest that the sensation of itchiness has a physiological role in risk estimation for scratched sites by translating the normally unrecognized risk into a decreased probability of successful itch-extinguishing and embodying it by the number of repeats of scratching bouts. Since the probability of successful itch-extinguishing is simply calculated using the maximum likelihood estimation method, it may be useful both in experimental scenes and clinical practices as an objective method to describe the quality of itchiness. The methodology of focusing on the number of repeats, as in this study, could also be applied to measure other physiological phenomena and diseases, such as the quality of coughing in asthma or the quality of repetitive behaviors in anxiety disorders.

## Results

We focused on the temporal intervals between scratching bouts in itch episodes. A single scratching bout is defined as a sequence of raising and lowering the hind leg, including scratching continuing over 150 ms in the SCLABA-Real system, which automatically records scratching bouts. However, an itch episode is not defined yet, and we thought that a sequence of bouts separated by very short inter-bout intervals seemed to express an itch episode (Figure 1A, underlines). We then established the hypothesis from this data that itch episodes consisted of the patterns of repeat of scratching bouts for duration of the emotion of itch (Figure 1B), and itch episodes and the number of scratching bouts were presented by a relational expression shown in Materials and Methods. All data of scratching bouts in the mouse are obtained as a CSV file (Figure 1C), and we can analyze the temporal intervals by calculating the difference between onset times (Figure 1D) and classify scratching bouts by inter-bout intervals into simplified (for example, binary) data (Figure 1E).

**Figure 1.**
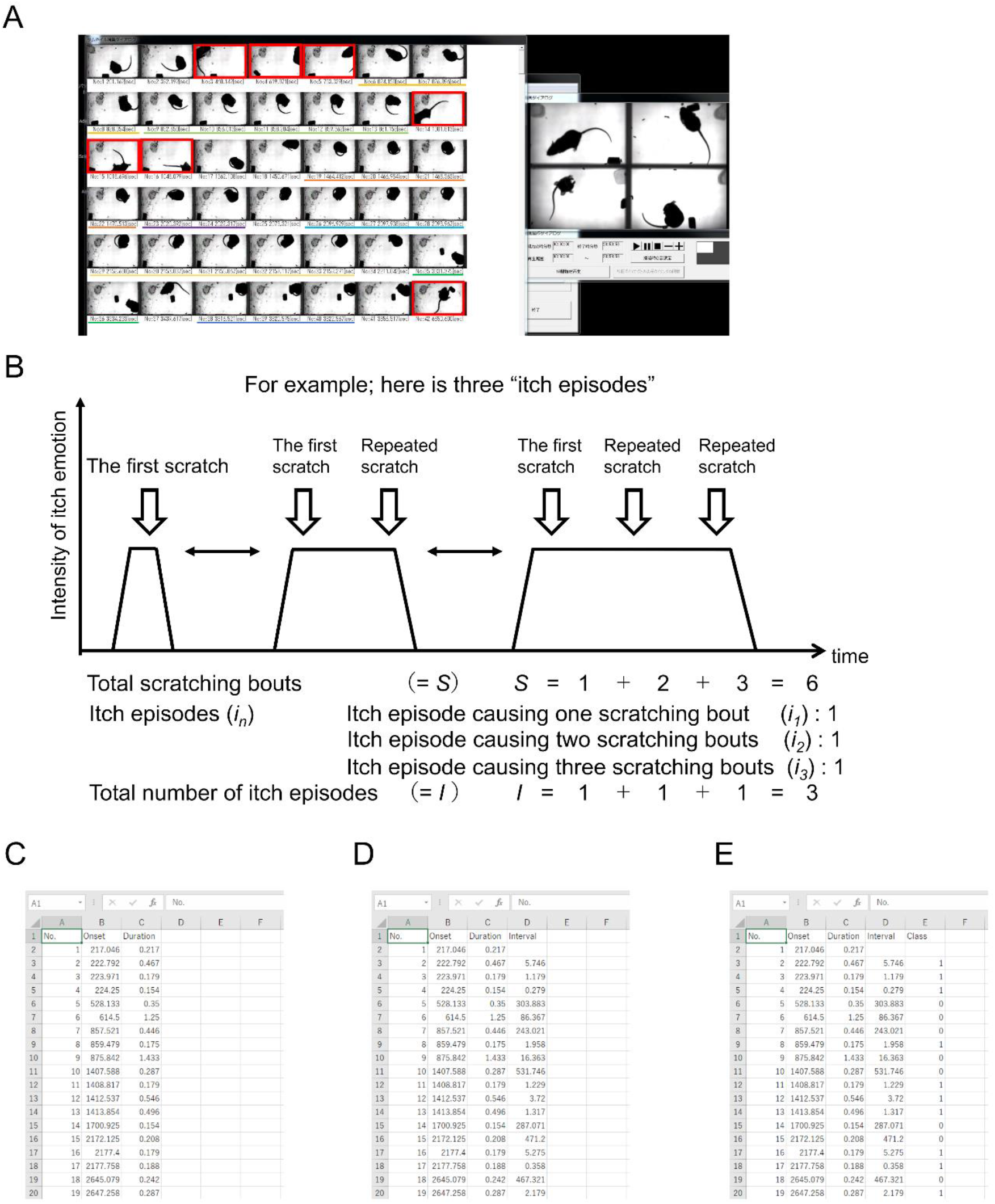
Hypothesized relationship between scratching bouts and itch episodes (A) Analysis of scratching bouts using the SCLABA-REAL video monitoring and analysis system. Small windows on the left show video segments in which a scratching bout was detected and manually verified (right). The time of occurrence of the bout is indicated under the window. The investigator underlined a sequence of windows with the same color to represent an episode of repetitive bouts separated by very short interruptions, thought constitute a single itch episode experienced by the mouse. The investigator marked some windows where the automatic segmentation algorithm erroneously included non-scratching bouts (including jump, dash) with a red box. (B) Hypothetical sequence of three scratching episodes of repetitive bouts, each assumed to correspond to an itch sensation. Trapezoids indicate the duration of the episodes, and the downward arrows indicate the onset of bouts. The episodes were classed by the number of bouts they consisted of (1, 2, 3, etc.), and the occurrence of episodes corresponding to each bout-number was counted (C) Example of a csv data file exported from the SCLABA-REAL system from a single recording session. Each row corresponds to a single bout; ordinal number, onset time, and duration are listed by the first, second, and third columns, respectively. Results of subsequent steps of analysis were saved in the same format. (D) Inter-bout time intervals. (E) Scratching bouts, classified either as the first in an episode (0) or a repeated one (1) within the criterion of time window (10 s).

A total of approximately 5,000 scratching bouts were obtained in 16 male and 16 female mice, each continually recorded in 24-h free-moving behavioral monitoring sessions (Supplemental Figure 1). The scratching bouts were classified according to the immediately preceding inter-bout time interval, as shown in Figure 1D. The inter-bout time interval varied considerably, but the majority (approximately 56%) was less than 10 s interval (Figure 2A). We defined the scratching bout over 10 s interval as the first scratching bout and the scratching bout under 10 s interval as the repeated scratching bout for a duration of the emotion of itch. In the data file of scratching bouts, the first or the repeated scratching bout was replaced by 0 or 1, respectively (Figure 1E).

**Figure 2.**
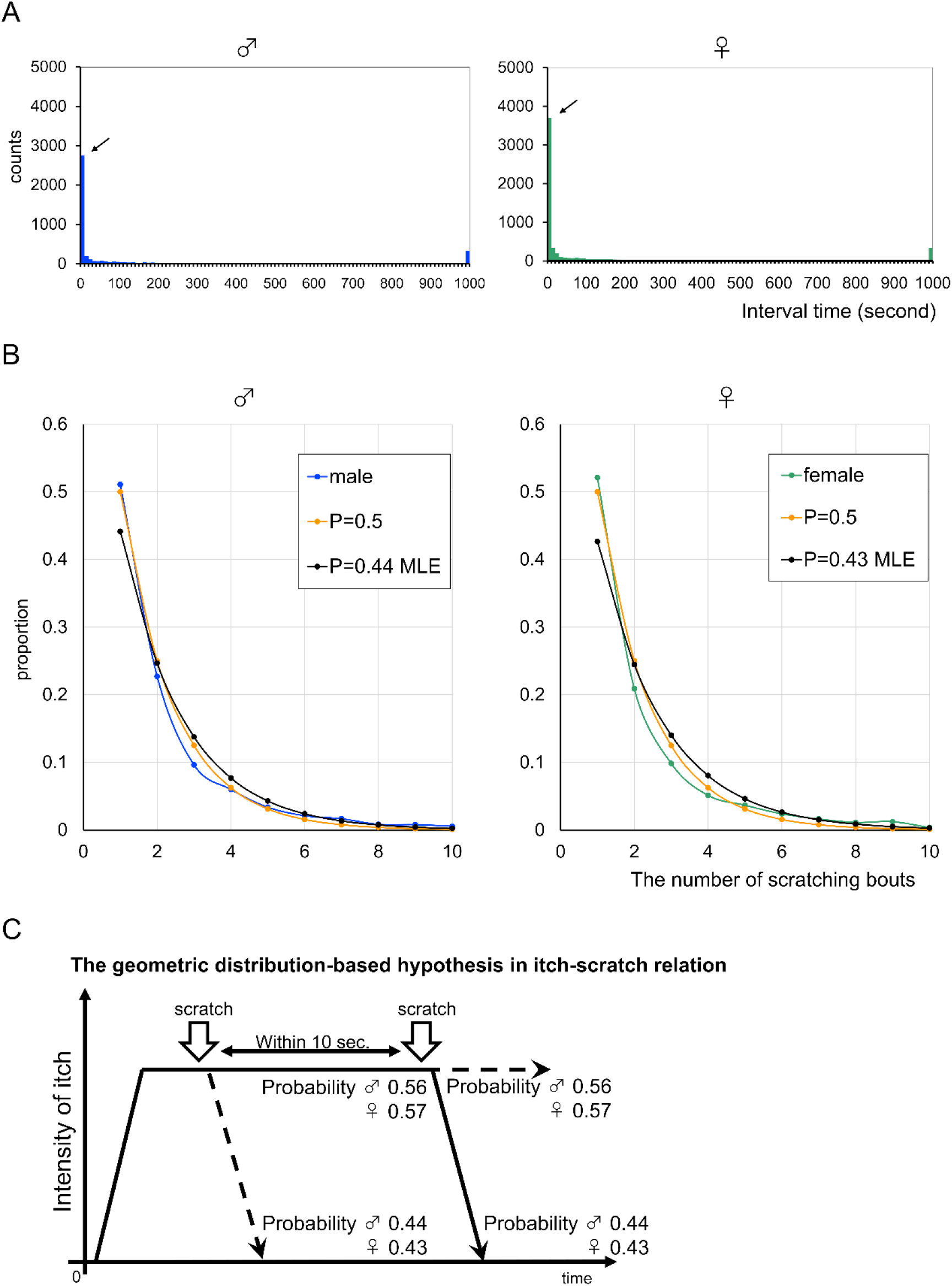
Patterns of the number of repeats of scratching bouts in healthy mice with back shaved (A) Scratching bouts grouped by their interval time. Over half of scratching bouts were under 10 s of interval in male and female mice (arrows). (B) Distribution of occurrence proportion of itch episodes causing a certain number of scratching bouts X axis indicated the number of scratching bouts in duration of the emotion of itch. Y axis indicated each proportion of itch episodes. Blue and green lines showed the distributions of male and female mice, respectively. Orange lines were described as 0.5 to the power of n, by analogy with the proportion of single scratching only. Black lines were obtained from the geometric distribution with success probabilities calculated by maximum likelihood estimation using actual data of male and female mice. P in the boxes means probability. MLE in the boxes means that maximum likelihood estimation was used. (C) New hypothesis of the relation between itch and scratching. These results indicated that scratching bouts were the Bernoulli trials until the emotion of itch was eliminated. The number of repeats of scratching bouts in an itch episode was controlled by the success probability. Probability calculated by maximum likelihood estimation was used in this figure.

According to this criterion, the patterns of the number of repeats of scratching bouts in male and female mice during the duration of itch emotion were represented by histograms (Supplemental Figure 2A, B). These were similar in shape each other and there was likely a same law in the pattern of repeats of scratching bouts. Summarizing the pattern data from 16 male and 16 female mice, we found that these patterns described smooth curves of decreasing and they were not normal distributions (Supplemental Figure 2C). Since it was difficult to infer the probability distribution from the measured values, we calculated the percentage of occurrences for the number of each repeat (1 (no repeat), 2, 3, …) (Figure 2B, Supplemental Figure 2D). Surprisingly, the distributions of occurrence percentages were almost same between male and female mice, and the distributions of the percentage values was close to (0.5)^1^, (0.5)^2^, (0.5)^3^, …, i.e., 0.5 to the power of n (n indicated the number of times of scratching bouts). To explain the interesting behavioral phenotype in mice, we considered the geometric distribution the best way possible. The geometric distribution is known to provide the distribution of the number of repeated trials until first success, which are independent Bernoulli trials with constant probability (of success). Because the end of the repeat of scratching bouts equaled to the successful elimination of the discomfort of itchiness, the probability deciding the fate of the number of repeats of scratching bouts was named as the probability of successful itch-extinguishing (SIE) in the present study (Figure 2C). When the distributions of the patterns of repeats of scratching bouts in male and female mice were applied to the geometric distribution, the probability of SIE were calculated to be 0.44 for males and 0.43 for females by maximum likelihood estimation (Figure 2B, line of MLE).

We found that healthy mice unconsciously repeated scratching bouts with constant success probability in spontaneous itch-scratch. However, it had been veiled whether itchy mice exposed by external factors also repeated scratching bouts obeying the geometric distribution. We then applied same methodology on model mice of itchy disease. Dry skin is common local skin disease with itchy all over the world and is caused by barrier destruction of stratum corneum. Dry skin of back was induced by treatment of mixture of acetone and diethyl ether following water in the same mice used in the study described above, and their scratching bouts were recorded for 24 hours (Supplemental Figure 3). Approximately 15,000 scratching bout data obtained from 24-hour behavioral records of 16 male and 12 female mice (data of 4 female mice was not saved because of recording PC crash) were classified by the time interval (Figure3A). More than 70% of scratching bouts was grouped into under 10-s interval, and it suggested that the repeat of scratching bouts increased after dry skin induction. Summarizing the pattern data from 16 male and 12 female mice with dry skin (Supplemental Figure 4A), these patterns also described smooth curves of decreasing and the probability of SIE appeared to be lower than 0.5 (Figure 3B, Supplemental Figure 4B). When the patterns of male and female mice fit manually the geometric distribution, 0.28 for male and 0.35 for female were better probabilities of SIE. Maximum likelihood estimation by using all patterns of repeat of scratching bouts resulted in 0.21 for males and 0.27 for females. These results indicated that scratching bouts were independent Bernoulli trials until the emotion of itch was eliminated, but the probability of SIE was changeable reflected on the skin condition (Figure 3C).

**Figure 3.**
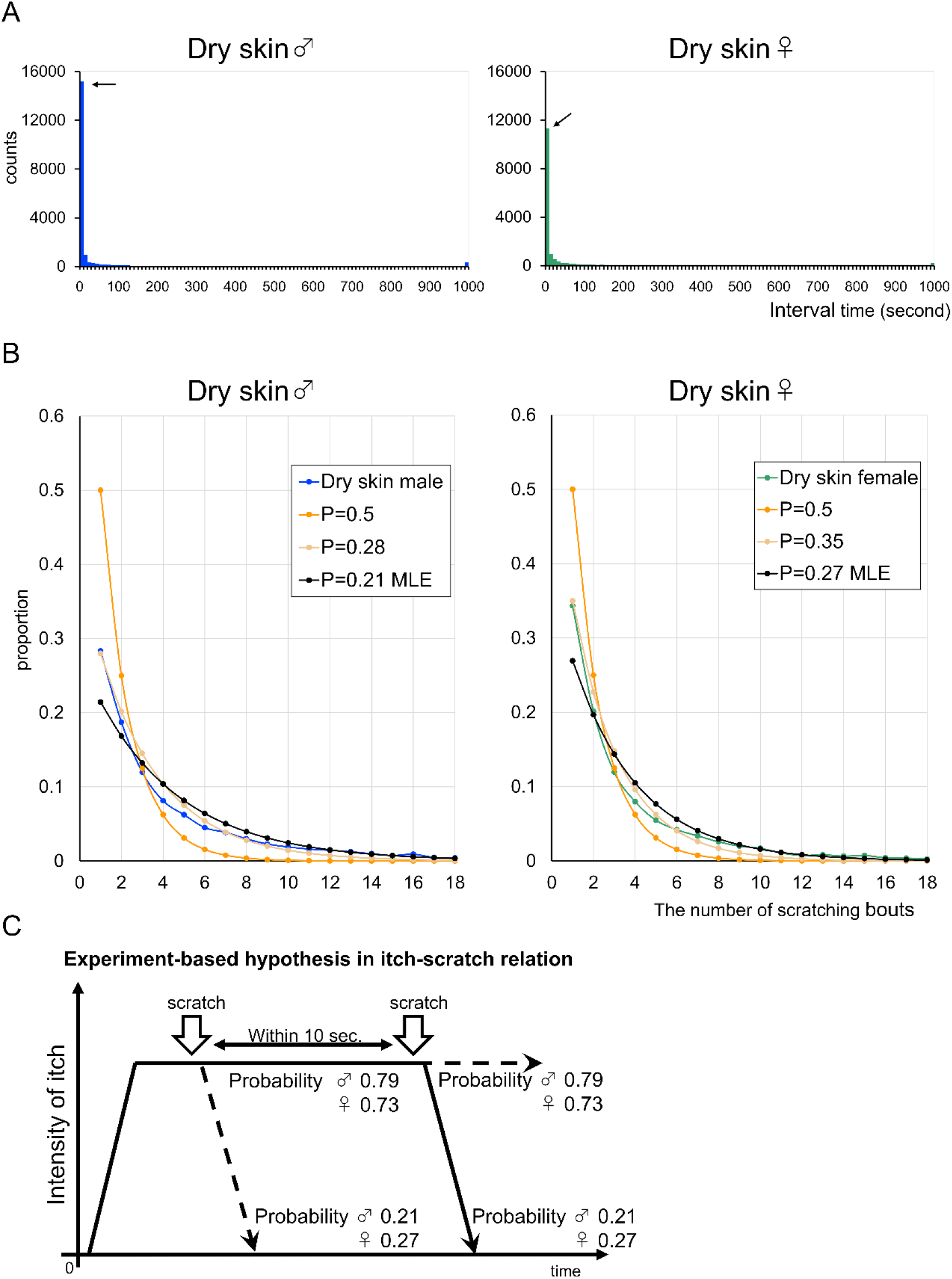
Patterns of the number of repeats of scratching bouts in mice with experimental dry skin induced in shaved back areas (A) Scratching bouts grouped by their interval time. Scratching bouts of dry skin mice of both male and female under 10-s interval time increased and the proportion was > 70% of all scratching bouts (arrows). (B) Distribution of occurrence proportion of itch episodes causing a certain number of scratching bouts. Lines obtained from experiment of dry skin mice indicated that their success probabilities decreased than that of healthy mice. Blue and green lines showed the distributions of male and female mice with dry skin, respectively. Orange lines were the geometric distribution with success probability 0.5, referring to Figure 2B. Beige lines showed the geometric distribution with success probability 0.28, by analogy with the proportion of single scratching only. Black lines were obtained from the geometric distribution with success probabilities calculated by maximum likelihood estimation using actual data of male and female mice with dry skin. P in the boxes means probability. MLE in the boxes means that maximum likelihood estimation was used. (C) Hypothesis of the relation between itch and scratching in dry skin. The probability of SIE decreased and the scratching bout tend to be easily repeated.

**Figure 4.**
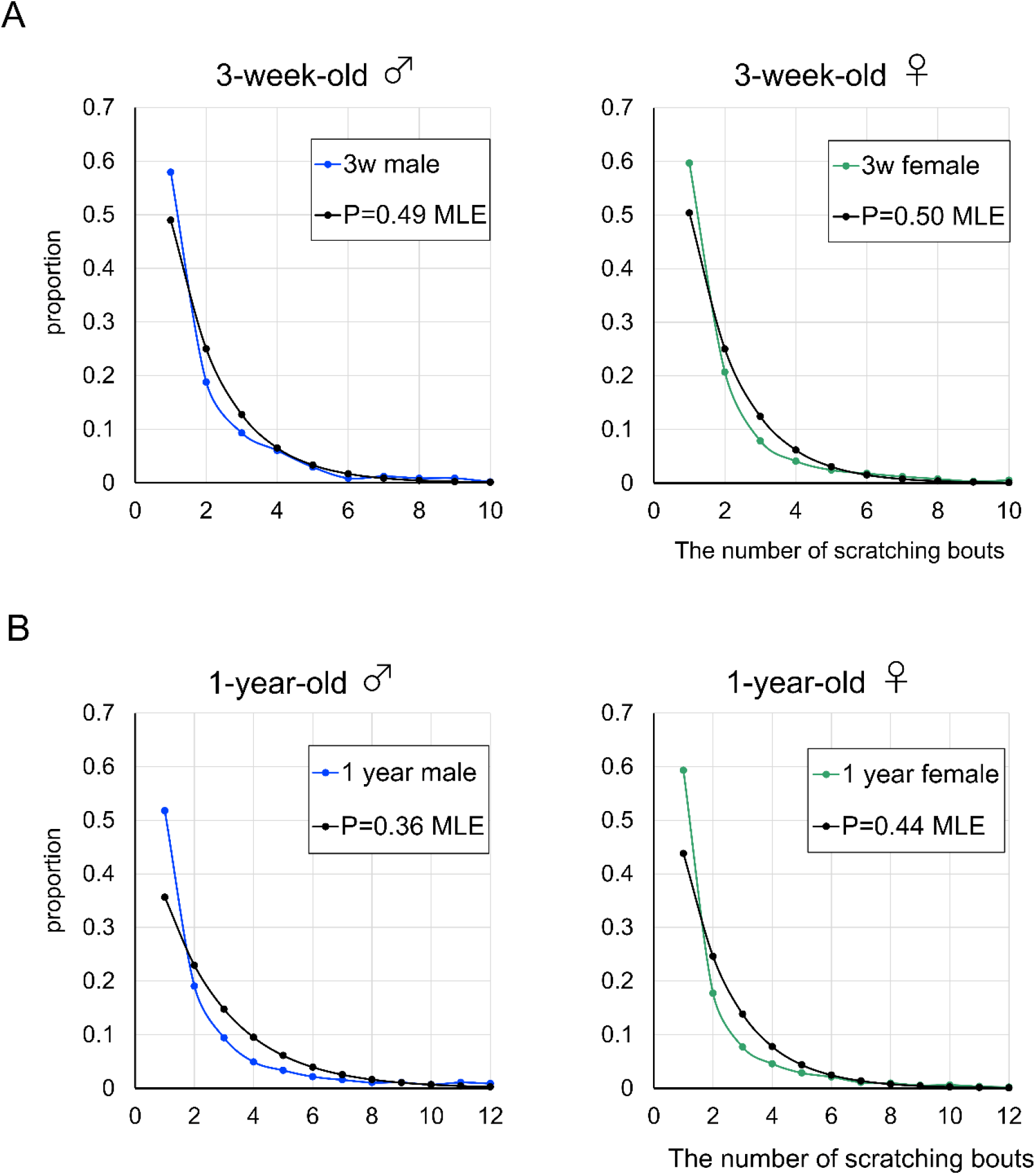
Patterns of the number of repeats of scratching bouts in mice raised from 3-week-old to 1-year-old Twelve of 3-week-old mice of both male and female without shaving were raised and the patterns of the repeats were investigated. (A) Distribution of occurrence proportion of itch episodes causing a certain number of scratching bouts in 3-week-old mice of male and female. Blue and green lines showed the distribution of the pattern of occurrence of itch episodes in male and female, respectively. Black lines showed the geometric distribution with success probability calculated by maximum likelihood estimation. These results indicated that the initial probability of SIE was innately set at 0.5. (B) Distribution of the occurrence proportion in 1-year-old mice. Blue and green lines showed the distribution in male and female, respectively. Black lines showed the geometric distribution with success probability calculated by maximum likelihood estimation. The probability of SIE in 1-year-old mice was decreased compared to that of 3-week-old mice, but the degree of decrease was smaller than that of dry skin mice.

Our results suggest that the probability of SIE determines the fate of the number of scratching bouts. Next, we investigated whether this is an innate or acquired ability during development. Twelve male and twelve female 3-week-old mice were raised for 1 year, and the probability of SIE was analyzed by setting a time point. When the patterns of repeat of scratching bouts in the 3-week-old mice was summarized, the distribution formed smooth curves of decreasing and it was not normal distributions (Figure 4A). Assuming the geometric distribution, the probability of SIE was 0.49 for males and 0.50 for females by maximum likelihood estimation. 1-year-old mice also displayed the distribution of the decreasing curves about the patterns of repeat of scratching bouts, but the values were 0.36 for males and 0.44 for females (Figure 4B). These results suggested that the probability of SIE is an innate ability rather than it acquired during development, and the probability of SIE decreased in old.

We claim that the repeat of scratching bouts is determined by constant probability-based decision making. When animals solve a problem, they usually use their learning ability to increase the probability of success and improve efficiency. Neural mechanisms that produce a constant probability of success for obtaining statistical data have never been envisioned therefore. In addition, the fact that weaning mice had already acquired the probability of SIE suggests that the probability of SIE is formed by innate simple neural circuits rather than by the development of complex neural circuits through learning. On-cell, Off-cell, and neutral cell suggest a probable relationship of mutual inhibition in pain (Miki K, 2002). We hypothesized that simultaneous excitation of mutually inhibitory neurons, followed by convergence to the excitation of one of the neurons, would produce two outcomes with the same probability each time (Figure 5A). This model was simple and agreed with the model of mutual inhibition of lateral inhibition reported by Koyama and Pujala (Koyama & Pujala, 2018). Based on the results of this study, it is likely that the scratching bouts is not targeted at the actual parasite, but rather at a fictional parasite. In other words, neurons that suggest the absence of a parasite and neurons that suggest the presence of a parasite are mutually inhibitory neurons because they represent mutually contradictory states. Stimulation by scratching bouts would result in the simultaneous excitation of these two neurons, resulting in a probabilistic convergence to a state where the imaginary parasite is either present or absent from the skin. Simple simulation of the interaction of mutually inhibitory neurons in case of homogeneous and heterogeneous (one was set at 10% stronger in inhibitory power) showed the probability at 0.49 and 0.27, respectively, calculated by maximum likelihood estimation method (Figure 5B). The process of calculation by two mutual neurons was shown in a movie.

**Figure 5.**
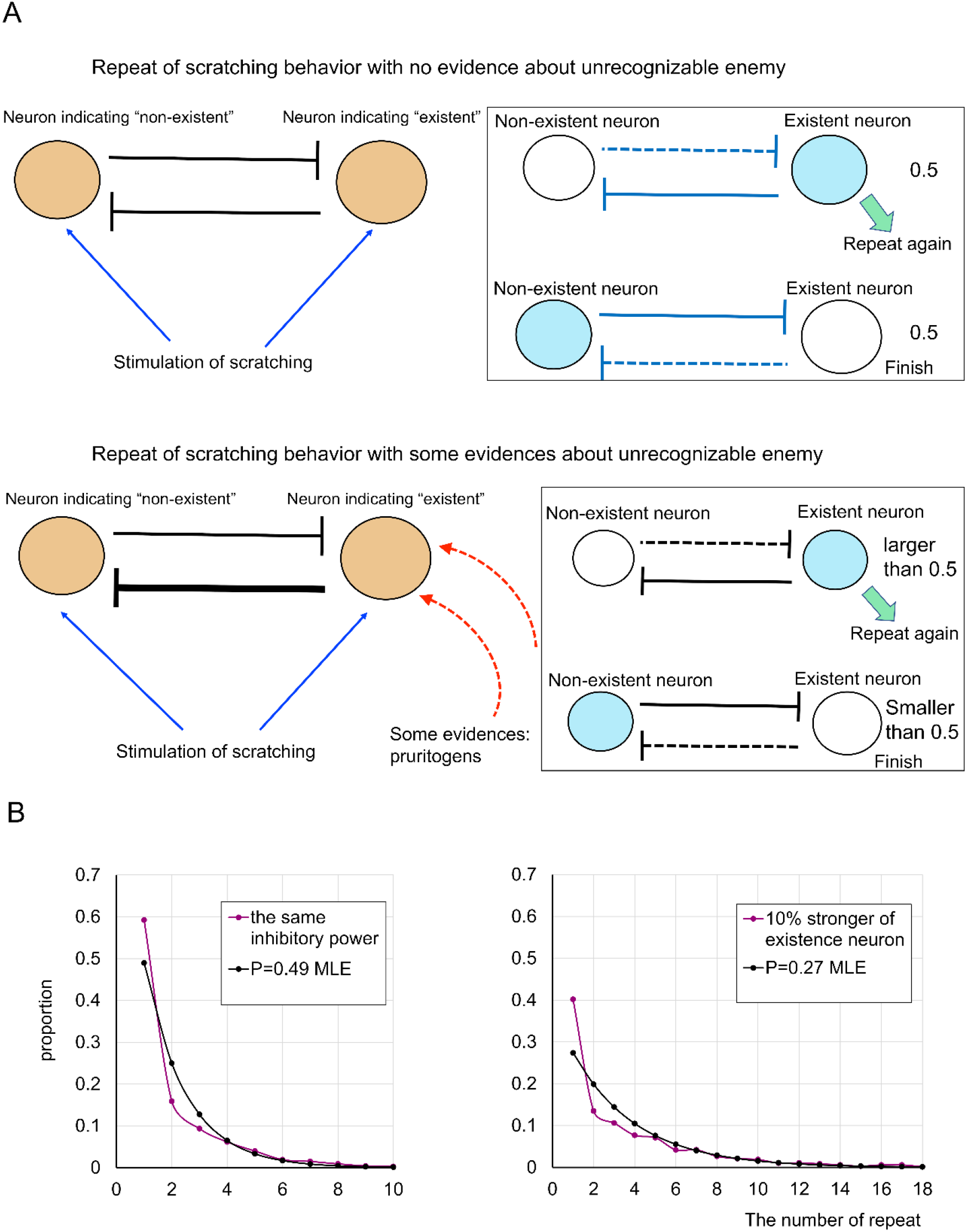
Model of probabilistic repeat by computing of simultaneous excitation of mutual inhibitory neurons The nature of probability is still unknown, but we speculated the mechanism of generation of probability in neural activity. For the struggle for existence of lives, to recognize the existence of enemy is the most important. However, lives are only capable of realizing enemies via their own senses, for example five senses in humans. Itch and the probability of SIE is the clue or the evidence of existence of unrecognized enemies. (A, above) Neurons indicating “non-existent” and “existent” were simultaneously excited by stimulation of scratching, and then the dual excitation converges to either neuron remaining firing. This process is performed regardless of the real existence of enemies such as parasites, and is repeated until existent neuron was suppressed. (A, lower) When real enemies invade the skin, the information of pruritogens transmits into the existent neuron and its inhibitory power increases by, for example, increasing the synaptic connection of mutual inhibitory neuron. Stimulation of scratching bouts evokes simultaneous excitation of mutual inhibitory neurons, but the converging result tends to remain exciting of existent neuron. (B) Results of the simulation of mutual inhibitory neurons. Left; In case that the same inhibitory neurons suppress reciprocally, the pattern of distribution of the number of repeats until converging an excitation of the non-existent neuron displayed that the probability of SIE was 0.49 calculated by maximum likelihood estimation. Right; Heterogeneous mutual inhibition that existent neuron has 10% stronger inhibitory power showed the lower probability of SIE at 0.27, calculated by maximum likelihood estimation.

### Movie

Movie of the model of probabilistic repeat by computing of simultaneous excitation of mutual inhibitory neurons

Left and right circles indicated non-existent and existent neurons, respectively. Excitation was expressed by rhythmic flashing. The fictional scratching bout added +1 to the count of scratch, and excite the two neurons. The converging to only existent neuron exciting evoked repeating the fictional scratching again.

## Discussion

The presence of parasites on itchy skin was first discovered in the 1830s (History of the Itch, 1870). Since that time, however, questions had arisen about the ability of such tiny, invisible organisms to cause itch. Although it is now clear that histamine and various pruritic substances cause itching, the possibility that parasitic organisms are the target of scratching bouts has not been ruled out. Many people know empirically that humans and other animals repeatedly engage in scratching bouts, but the pattern of repeated behavior has not been scientifically examined. We conducted a study based on the hypothesis that the repeated pattern of scratching bouts represents the quality of itchiness. As a result, we found that the repeated pattern of scratching bouts follows the geometric distribution, which means that the repeated trials are based on a constant probability. This result provided strong evidence that the purpose of the scratching bouts is not to target actual parasites in the skin. At the same time, it is surprising that a constant probability of success is set for the scratching behavior, which was thought to be repeated unconsciously. As the simple mechanism of probability generation in the central nervous system, we proposed a probability control model based on the simultaneous excitation of neurons in a mutually inhibitory relationship. In this model, the probability of SIE is structurally fixed according to whether the inhibitory ability of two neurons is the same or different, and can be embodied by repeated scratching bouts, which resembles the way information is saved in a quantum memory. The details are still unclear, and more research is needed to understand how neurons can express constant probabilities.

If the number of times that scratching bouts are repeated is based on a strict law, scratching behavior must be a physiologically and biologically meaningful behavior. For example, skin barrier disruption by 4% SDS application induced migration of regulatory T cells (Toyama et al., 2021). Epidermal nerve fibers increased transiently by skin barrier disruption following acetone treatment (Tominaga et al., 2007). These findings indicate that disruption of the skin barrier enables us to take care for ourselves. Because healthy mice experience the probability of SIE at 0.5 from 3 weeks of age, it is likely that the subjective success probability is also approximately 0.5. Because a subjective probability of success of 0.5 in humans represents the maximum motivation (Atkinson, 1957), it is unsurprising that the probability of SIE is maintained at 0.5 in humans and other animals. When the probability of SIE is reduced in mice with dry skin, it will be possible to identify the skin area at risk only when mice recognize that the probability of SIE is reduced by actual scratching bouts. The calculation of the reward prediction error may play an important role in recognizing the decrease in the probability of SIE (Eshel et al., 2015). The lattice structure like Ising model of micro columns including somatostatin-expressing and parvalbumin-expressing neurons may have a role in the probability of SIE because synchronistic activation in the same micro column is evoked by the input of the same excitatory neuron in the somatosensory cortex (Maruoka H, 2017). To test whether the risk estimation role of itch is important for the animals, it is necessary to breed mice that do not feel itchy, such as GRPR-deficient mice, in a conventional environment for a long time (Pagani et al., 2019). Although various kinds of pruritogens cause both itch and pain in almost same qualia (Klein et al., 2021), there may be some differences in the effects on the probability of SIE.

In this study, the emotion of itch evoked by dry skin was investigated as a typical symptom of itchy skin. It is necessary to analyze itchiness caused by other etiologies such as atopic dermatitis. If there is a distinct itch in which the pattern of repeated scratching bouts does not form a geometric distribution, it will be easy to infer the disease causing the itch from that pattern and will be useful as a noninvasive and nonverbal diagnosis. In addition, if some patients complain of itch even if the probability of SIE is not reduced, this may indicate a psychological effect of anxiety or addictive scratching that is greater than the actual risk (Ishiuji, 2019). The behavioral study of itch in animals, including humans, requires further experimentation.

The use of the probability of SIE in itch research may provide another way to evaluate the efficacy of antipruritic drugs, which is not clear from examining the number of scratching bouts. The number of scratching bouts has been used as a measure of itch quality in all previous studies of itch, but differences between males and females have been reported (Stumpf et al., 2013; Takanami et al., 2021). Focusing on the number of scratching bouts in our 24-h recording data, we found that the number of scratching bouts per hour varied from 0 to 99 in healthy mice (Supplemental Figure 1A). The pattern of the number of scratching bouts per hour has not been determined yet, but the number of scratching bouts per hour in the light period was less than that in the dark period and the average number of scratching bouts in the dark period was higher in females than in males (Supplemental Figure 1B). Data on the number of scratching bouts in bright and dark environments were then incomparable. The total number of scratching bouts and scratching time over 24 h was higher in females, but the scratching time per scratching bout was similar in males and females (Supplemental Figure 1C-E). This suggested that there was a difference in the number of itch episodes. The number of scratching bouts in the dark period was significantly higher in females than in males, but there was no difference in the light period. The rate of increase of scratching bouts in the dark period relative to the light period was approximately 1.8 times in both males and females, with no significant difference (Supplemental Figure 1F-H). This suggested that the frequency of itch episodes increased to the same extent in both males and females in the dark period. In a comparison of the number of scratching bouts in the light and dark periods in males and females, the number of scratching bouts was significantly higher in the dark period (Supplemental Figure 1I, J). We examined the hypersensitivity to itch sensation evoked by mechanical stimuli among time periods when the number of scratching bouts in males and females was or was not significantly different (Supplemental Figure 1B, asterisks) and found no significant difference in all conditions (Supplemental Figure 1K-N). This suggested that the change in the number of scratching bouts in the light and dark periods was not related to the hypersensitivity to physical stimuli. In mice induced with dry skin, the number of scratching bouts increased to >50 per hour, with a maximum of >400 per hour (Supplemental Figure 3A). The average number of scratching bouts per hour was similar in the light and dark periods (Supplemental Figure 3B). There was no significant difference in the number of scratching bouts over 24 h between males and females, but the number of itch episodes was significantly higher in females (Supplemental Figure 3C). This suggested that the decrease in the probability of SIE was greater in males. In a comparison of the number of scratching bouts of males and females in the light and dark periods, the number of scratching bouts in the light period was significantly higher in males, whereas there was no significant difference in females (Supplemental Figure 3D). We examined the correlation between the number of scratching bouts and the number of itch episodes under normal conditions and after dry skin induction (Supplemental Figure 3E). In males, there was a positive correlation between the number of scratching bouts and the number of itch episodes in the healthy state after dry skin induction; however, these indices were not positively correlated in females. These results suggested that many conditions must be satisfied when comparing males and females in terms of the number of scratching bouts, and that it will be difficult to conduct itch studies in females. The probability of SIE was a common feature in healthy males and females. The index will be useful for comparing the quality of itchiness between the sexes.

### Limitation

#### Definition of the interval between itch episodes

The definition of an itch episode may be controversial. In this study, we inferred itchiness in mice from the perspective of “itchy because scratching was observed” rather than “itchy because itch emotion was observed.” The design of our research was not based on the James–Lange theory or Cannon–Bard theory about the origin of emotion. Humans do not always engage in scratching bouts when they experience itchiness, but they engage in scratching bouts even if causes skin damage. This suggests that there is a gap between the persistence of itch and the execution of the scratching behavior, with or without emotion. In other words, the attentional duration in itching corresponds to the interval of a scratching bout within an itch episode, and the emotional duration in itching corresponds to the duration of scratching bouts. Because animals, except humans, cannot report itchy, it is extremely difficult to define an itch episode completely and correctly. In a study of mechanical alloknesis, which refers to hypersensitivity causing scratching bouts induced by light touch stimuli, scratching bouts that occurs within 10 s after physical stimulation to the skin were defined as responsive bouts to the stimulus (Chen et al., 2020). This implies that the attentional duration of light touch stimuli was assumed to be up to 10 s before the emotion of itch was aroused. In addition, the researcher in this study had an empirical intuition that mice often started moving >10 s after the end of one scratching bout. Furthermore, considering the statistical data on the interval of scratching bouts shown in Figure 2A, the definition of the interval between itch episodes was delimited by 10 s. Although there might be a pattern of repeated scratching bouts with attention durations of >10 s, the statistical data suggested that such cases, if any, were few in comparison to the total and would not have a significant impact on the results of our analysis.

## Materials and Methods

### Animals

Care and handling of all animals conformed to the NIH guidelines for animal research, and all animal procedures were approved by the Institutional Animal Care and Use Committee of Juntendo University Graduate School of Medicine. Seven-week-old C57BL6/J male (n = 16) and female (n = 16) and 3-week-old C57BL6/J male (n = 12) and female (n = 12) mice supplied by Charles River (Japan) were maintained in the experimental animal facility of Juntendo University Graduate School of Medicine under a 12:12-hour light: dark cycle at 22–24°C, with food and tap water provided *ad libitum*. Mice at 3 weeks of age were maintained until they grew to 1 year of age.

### Experimental induction of dry skin

Dry skin was induced experimentally, as previously described (Miyamoto et al., 2002). The skin was shaved in the rostral back area at least 1 week prior to the experiment. To break the skin barrier, cotton patch (2 × 2 cm^2^) soaked in a 1:1 acetone/diethylether mixture was applied to the shaved skin area for 15 s, followed by a cotton patch soaked with distilled water for 30 s. The treatment was performed twice daily for 6 days.

### Automatic video recording of scratching bouts

Measurements were made for individual animals one-by-one in 24-h sessions. The animal was transferred into the experiment cage and its scratching bouts was continuously recorded by a real-time video monitoring and analysis system (SCLABA-Real system) over a 24-h period, as previously described (Orito, 2004). A researcher set the timer to start the session of automatic recording of scratching bouts 1h after leaving, and no one entered the room during the recording session. A single bout is a single uninterrupted sequence of periodic hind limb movements corresponding to one or more scratches, lasting for a minimum of 150 ms (the elapsed time roughly corresponding to a single scratch). Detection of scratching bouts and extraction of video segments of scratching were done by automatic analysis of the video records from the session. Segments were screened for false positive (e.g., walking, wiping, and jumping) and these were manually removed. The data thus preprocessed were used in further analysis, including frequency and duration of scratching.

### Counting of the number of scratching bouts for itch episodes

Video segments of scratching were collected as described above. The three parameters extracted for each bout included the ordinal number in the sequence of occurrence; the bout onset time, measured from the recording start time; and the bout duration. These were exported in a csv file organized by columns to be used as input for subsequent steps of analysis. Next, using a spread sheet (MS Excel), we derived the inter-bout time interval (the time elapsed between the onset times of subsequent bouts), and classified the bout either as a “first bout” (class 0) if the preceding inter-bout interval was longer than a criterion (10 s), or a “repeating bout” (class 1) otherwise. The criterion was set according to the result of Figure 2A and explained in the Discussion section.

Using the statistical analysis package R, the binary bout sequences were analyzed for the length of repeat sequences (defined by the length of uninterrupted sequence of 1’s). Itch episodes were defined by a bout sequence starting with 0 until (but not including) the next 0 event. A single bout episode was represented by the sequence [0]; an episode of two bouts, by [0,1], an episode of 3 bouts, by [0,1,1]; etc.

To determine the distribution of itch episodes binned by bout-count, we counted the number of itch episodes observed for each bout-count for each animal. For verification of the accuracy of the counting, we recalculated the number of scratching bouts from the number of itch episodes.

### Analysis of repetitive scratching

To determine the empirical probability distribution of itch episodes from the observed frequency distribution of itch episodes, we normalized episode counts in each bout-count bin by the total number of episodes. We chose the family of geometric distributions to model the probability distribution because it provides a nice conceptual explanation of repetitive scratching.

The geometric distribution of parameter *k* is the discrete probability distribution of the number of independent Bernoulli trials needed to get one success at the end. If the probability of success on each trial is *p*, then the probability that the *k*th trial (out of *k* trials) is the first success is

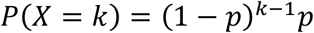

Conceptually, a bout of scratching could be thought of as probabilistic event of itch-extinguishing behavior, with each bout having probability *p* of extinguishing the itch sensation. Then the probability that itching is extinguished by the *k*th bout in an episode of repetitive scratching bouts is given by the geometric distribution of parameter *p*. We may call the success probability in this study the probability of SIE. By definition of *p* = *P*(*X* = 1), and the single-trial value of the empirical probability distribution was used as the initial value for the optimization. Maximum likelihood estimation was used to optimize the value of the single-trial probability of success, *p*, to fit the geometric distribution to the empirical distribution. The p value obtained by maximum of likelihood function for the geometric distribution was calculated as reciprocal number of the average of all itch episodes.

### Probabilistic control model by mutually inhibitory neurons

Simulations and video presentation of model data of mutually inhibitory neurons involved in controlling repetitive scratching bouts were made using custom code written with open-source software (Pure Data; version 0.51-4).

Three parameters were used to model a template inhibitory neuron, including the firing rate (initially set to constant 20 Hz), probability of successful firing (initially set to 99.5%), and inhibitory power per one firing (set to reduce the probability of successful firing by 5%). To visualize simulated neural firing with a “flash”, the bang object was used to trigger an instantaneous brief flip of the color from white to black in the display representing the neuron.

To simulate control of scratching bouts, two inhibitory neurons were cloned from the same template and set to mutually inhibit each other. One neuron represented the presence of an imaginary parasite; the other represented homeostasis by the elimination of the imaginary parasite. Simulation of scratching was started by the press of the “itch start” button that triggered the two neurons to begin firing and lasted until their firing converged. If firing converged to the neuron that signified the presence of imaginary parasites, the next bout of simulated scratching was triggered. The simulated scratching bouts repeated until successful convergence to the neuron that signify the elimination of imaginary parasites. The above sequence of scratching bouts constituted a single simulated trial.

The interaction dynamics of two mutual inhibitory neurons was captured in video using an Xbox game bar preinstalled on a personal computer running under the MS Windows operating system. Simulated scratching incidents were summarized in a spread sheet (MS Excel) and the probability of SIE was calculated by maximum likelihood estimation.

### Itch sensitivity test by mechanical stimuli of von Frey filaments

At least 1 hour before the start of the experiment, the mice were caged one by one and acclimatized. At 5:00 and 17:00, the researcher applied total 20 times of weak stimulation of von Frey filaments (0.07 and 0.16 g) to shaved back areas with >10 s intervals. The criteria of itch evoked by weak stimulation was whether scratching bouts started within 10 s after the weak stimulation. The number of the reaction was showed as alloknesis score.

### Statistical analysis

Students’ t-test with p < 0.05 significance was performed for the male-female comparison of the data in the supplemental figures. BellCurve for Excel (Social Survey Research Information Co., Ltd., Tokyo, Japan) was used for the regression analysis of frequency distribution of itch episodes by bout-count (Figure 2 and 3), and the scatter plot matrix in (Supplemental Figure 3-1).

## Acknowledgements

We would like to thank laboratory animal care janitors in Juntendo University and Enago (www.enago.jp) for the English language review. This work was supported by a President’s Grant for Interfaculty Collaboration, Juntendo University (2020-1) and a JSPS KAKENHI Grant Number JP20H03568.

## Competing interests

No competing interests were involved in this study.

---

**Supplemental Figure 1.**
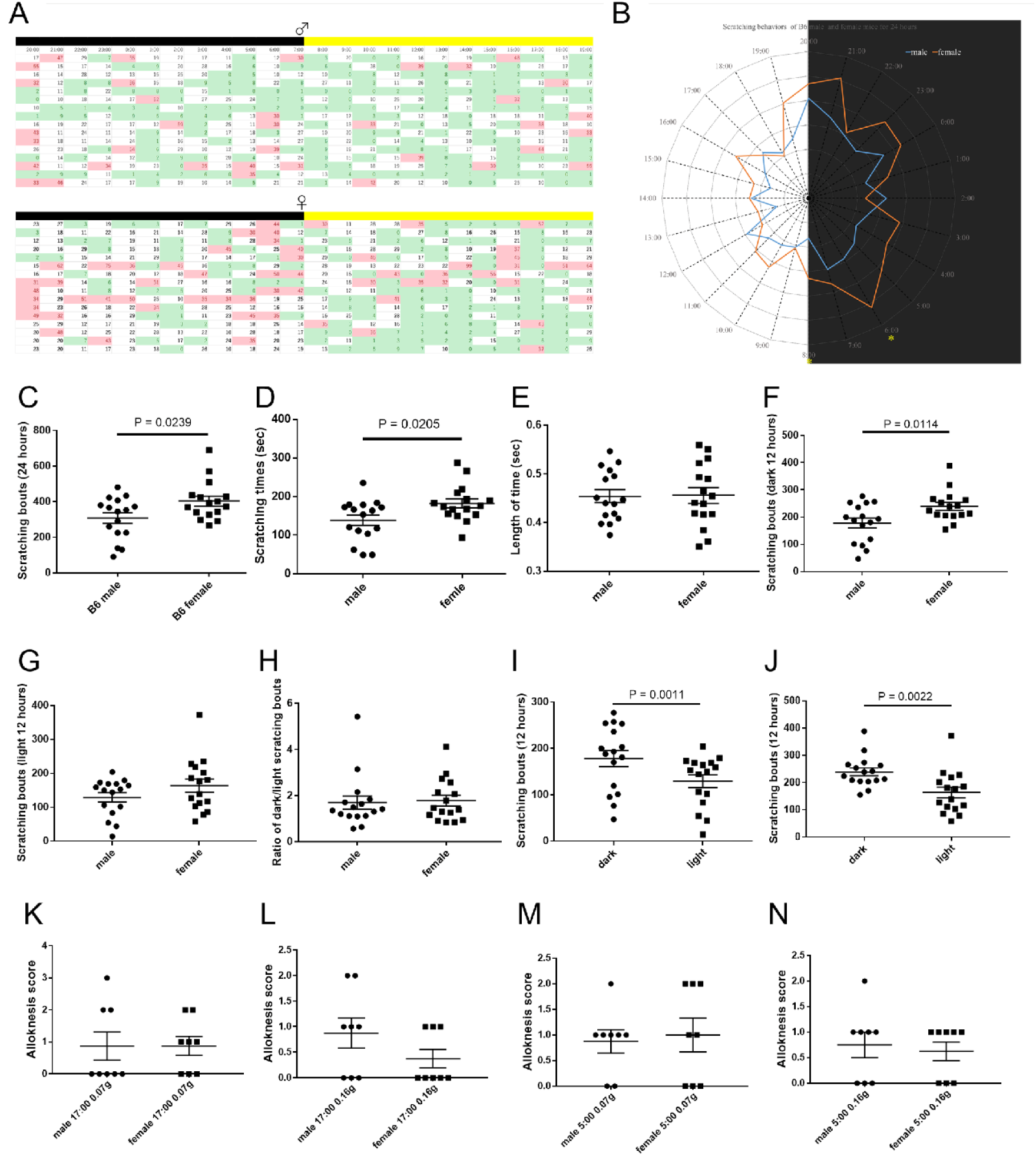
Statistical analysis of scratching bouts observed in non-treated mice during a 24-h recording session. (A) Hourly counts of recorded scratching bouts organized in a table, with each row for a different animal, in 16 male animals (top), and 16 females (bottom). Black (yellow) bars atop the tables represent the 12-h dark (light) periods. Boxes in white indicate near-median counts (10–29 range); green, low counts (<10); red, high counts (>30). (B) Polar plot of the circadian variation of the average hourly count in males (blue) and female (orange) animals. Asterisk marks significant (p < 0.05) male-female difference. (C) The total number of bouts, organized separately for the male (filled circles on the left) and female (filled squares on the right) group. Each symbol represents one animal. Three horizontal bars represent mean ± SEM. Bar indicates significant between-group differences (p < 0.05). Data in subsequent panels follow the same organization. (D) Total duration (s) of scratching bouts (summed across bouts). (E) Mean duration of bouts (ms). (F) Total number of bouts, restricted to the 12-h dark cycle. (G) Same as (F), for the light cycle. (H) Dark/light ratio of total counts. (I) Light-dark comparison of counts, in males. (J) Same as (I), for females. (K–N) Comparison of itch hypersensitivity of the sexes at different time zones and strengths of stimulus. Male vs female at 17:00 by 0.07 g (K) and 0.16 g filaments (L), and at 5:00 by 0.07 g (M) and 0.16 g filaments (N).

**Supplemental Figure 2.**
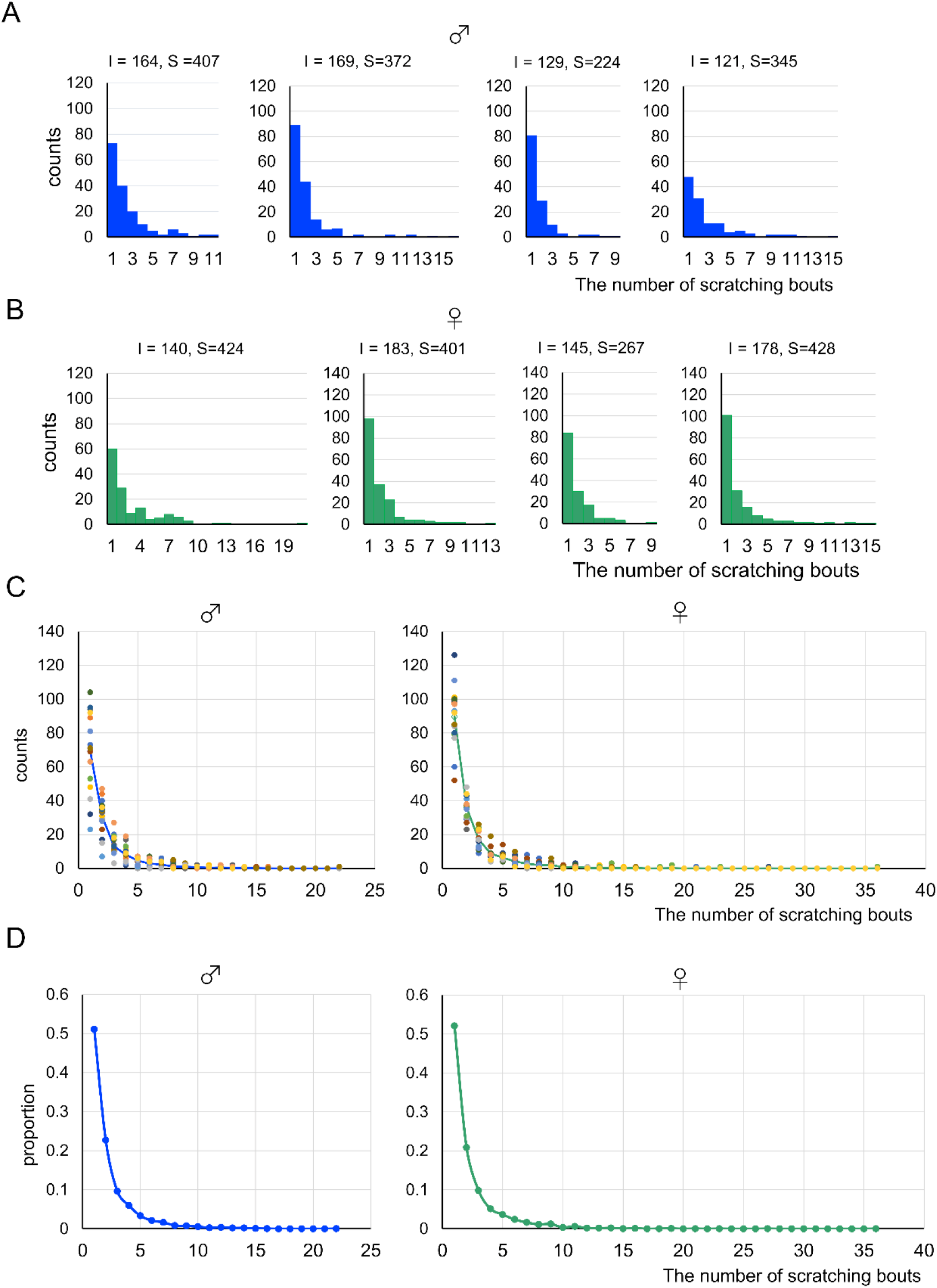
Frequency distribution of itch episodes by bout counts. Histograms plot the number of episodes (vertical axis) observed for increasing bout counts per episode (bins). Representative data from (a) 4 male mice and (B) 4 female animals. Notice the monotonically decreasing shape of the distributions. (C) Data from all 16 separately summarized for male (left) and female (right) animals. Data for each animal is plotted in distinct color. (D) Same data as in (C), expressed as fractions (summed across animals in each bin, and normalized by total number of episodes) to derive the discrete empirical probabilities. I: total number of episodes; S: total number of bouts.

**Supplemental Figure 3.**
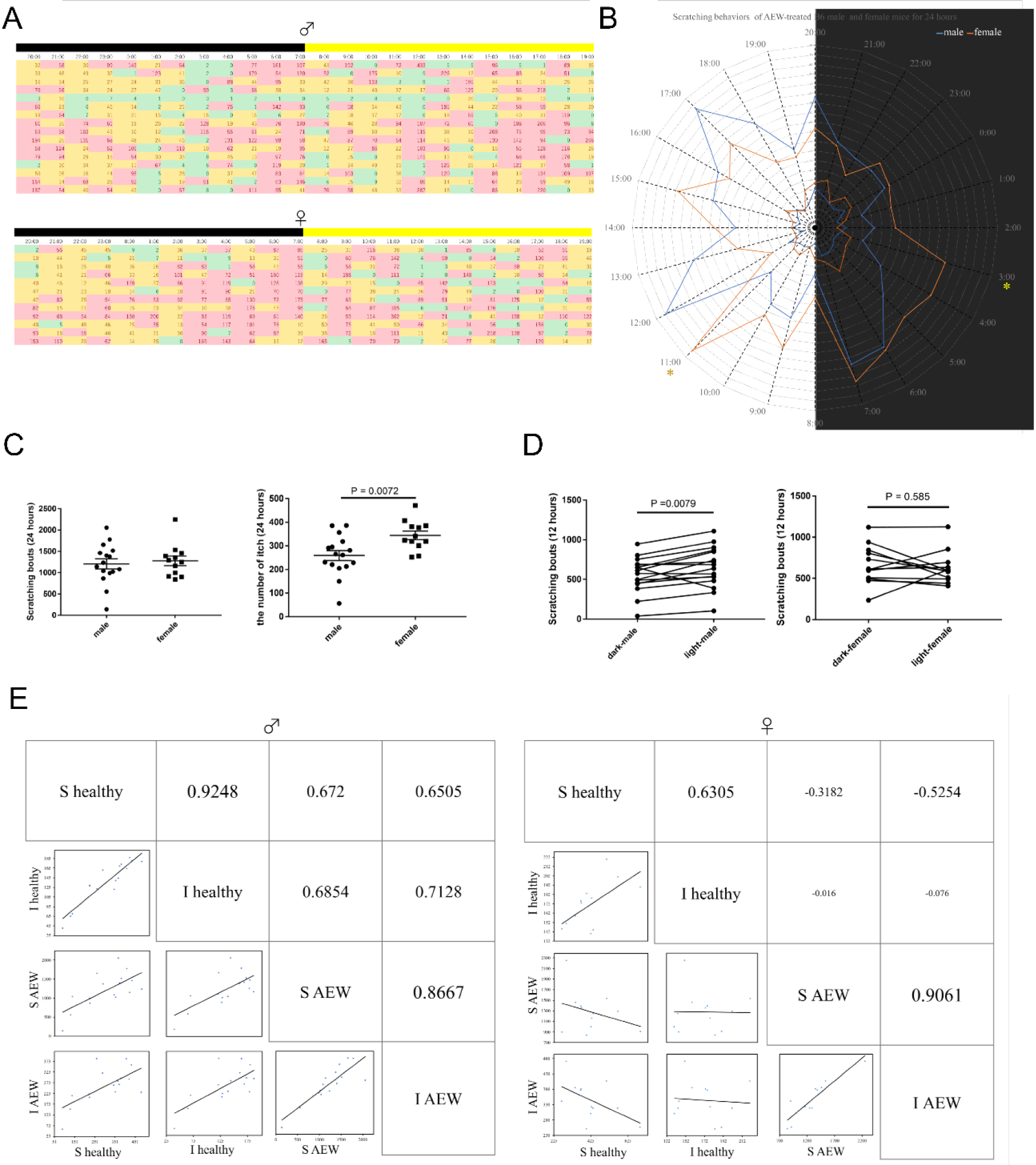
Analysis of the number of scratching bouts of dry skin mice for a day (A) The number of scratching bouts per an hour (columns) after dry skin induction was shown by male and female individuals (rows). The black/yellow zones indicated the light off/on periods respectively. Boxes colored by green, yellow, and red indicated fewer (<10), median (10~50), and larger (50<) scratching bouts in one hour. (B) Scratching averages after dry skin induction per an hour in male (blue) and female (orange) were shown by radar charts. To compare the number of scratching bouts before/after dry skin induction, radar chats before the induction as shown in Figure 1A were put inside. Asterisks mean significant differences at p < 0.05. (C) Comparing the number of scratching bouts (left) and itch episodes (right) for 24 h between male and female mice. (D) Comparing the number of scratching bouts between dark 12 h and light 12 h in males (left) and females (right). (E) The correlation matrix between the number of scratching bouts and itch episodes in male (left) and female (right) mice. Dots indicated mice individuals. S and I indicates “scratching bouts” and “itch episodes.” AEW indicates the status after dry skin induction following treatment with a mixture of acetone and diethyl ether and with water. Numbers in boxes indicate the values of the correlation coefficient.

**Supplemental Figure 4.**
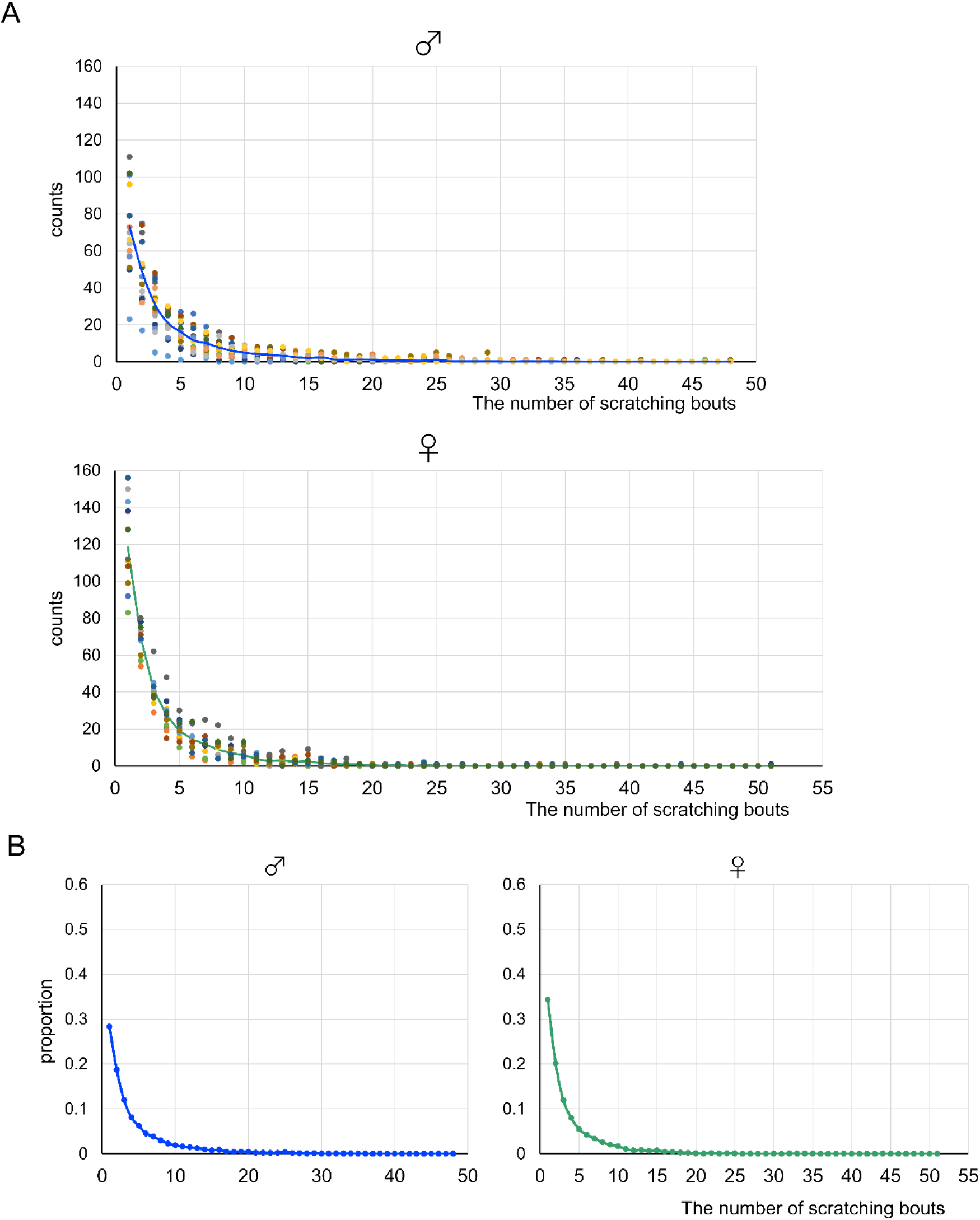
Patterns of occurrence of itch episodes in dry skin mice (A, B) Summary of the counts (A) and proportion (B) of itch episodes in 16 male (upper) and 12 female mice (lower, the data of 4 mice were broken by PC trouble). Dots colored indicated mice individuals.

## Notes

### Competing Interest Statement

The authors have declared no competing interest.

## References

Atkinson, J. W. (1957). Motivational determinants of risk-taking behavior. Psychol Review, 64(6), 359–372. https://doi.org/10.1037/h0043445

Chen, S., Gao, X. F., Zhou, Y., Liu, B. L., Liu, X. Y., Zhang, Y., Barry, D. M., Liu, K., Jiao, Y., Bardoni, R., Yu, W., & Chen, Z. F. (2020). A spinal neural circuitry for converting touch to itch sensation. Nat Commun, 11(1), 5074. https://doi.org/10.1038/s41467-020-18895-7

Eshel, N., Bukwich, M., Rao, V., Hemmelder, V., Tian, J., & Uchida, N. (2015). Arithmetic and local circuitry underlying dopamine prediction errors. Nature, 525(7568), 243–246. https://doi.org/10.1038/nature14855

Girolomoni, G., Luger, T., Nosbaum, A., Gruben, D., Romero, W., Llamado, L. J., & DiBonaventura, M. (2021). The Economic and Psychosocial Comorbidity Burden Among Adults with Moderate-to-Severe Atopic Dermatitis in Europe: Analysis of a Cross-Sectional Survey. Dermatol Ther (Heidelb), 11(1), 117–130. https://doi.org/10.1007/s13555-020-00459-8

Hart, B. L. (1990). Behavioral adaptations to pathogens and parasites: five strategies. Neuroscience and biobehavioral reviews, 14(3), 273–294. https://doi.org/10.1016/s0149-7634(05)80038-7

History of the Itch. (1870). Edinburgh medical journal, 15(8), 756–757.

Ishiuji, Y. (2019). Addiction and the itch-scratch cycle. What do they have in common? Exp Dermatol, 28(12), 1448–1454. https://doi.org/10.1111/exd.14029

Klein, A., Solinski, H. J., Malewicz, N. M., Ieong, H. F., Sypek, E. I., Shimada, S. G., Hartke, T. V., Wooten, M., Wu, G., Dong, X., Hoon, M. A., LaMotte, R. H., & Ringkamp, M. (2021). Pruriception and neuronal coding in nociceptor subtypes in human and nonhuman primates. Elife, 10. https://doi.org/10.7554/eLife.64506

Kogo, N., Kern, F. B., Nowotny, T., van Ee, R., van Wezel, R., & Aihara, T. (2021). Dynamics of a Mutual Inhibition Circuit between Pyramidal Neurons Compared to Human Perceptual Competition. J Neurosci, 41(6), 1251–1264. https://doi.org/10.1523/JNEUROSCI.2503-20.2020

Koyama, M., & Pujala, A. (2018). Mutual inhibition of lateral inhibition: a network motif for an elementary computation in the brain. Curr Opin Neurobiol, 49, 69–74. https://doi.org/10.1016/j.conb.2017.12.019

Maruoka H, N. N., Tsuruno S, Sakai S, Yoneda T, Hosoya T. (2017). Lattice system of functionally distinct cell types in the neocortex. Science, 358(6363), 610–615.

Miki K, Z. Q., Guo W, Guan Y, Terayama R, Dubner R, Ren K. (2002). Changes in gene expression and neuronal phenotype in brain stem pain modulatory circuitry after inflammation. J Neurophysiol, 87(2), 750–760. https://doi.org/10.1152/jn.00534.2001.

Misery, L., Dutray, S., Chastaing, M., Schollhammer, M., Consoli, S. G., & Consoli, S. M. (2018). Psychogenic itch. Transl Psychiatry, 8(1), 52. https://doi.org/10.1038/s41398-018-0097-7

Miyamoto, T., Nojima, H., Shinkado, T., Nakahashi, T., & Kuraishi, Y. (2002). Itch-associated response induced by experimental dry skin in mice. Jpn J Pharmacol, 88(3), 285–292. https://doi.org/10.1254/jjp.88.285

Mui, J. W., Willis, K. L., Hao, Z. Z., & Berkowitz, A. (2012). Distributions of active spinal cord neurons during swimming and scratching motor patterns. J Comp Physiol A Neuroethol Sens Neural Behav Physiol, 198(12), 877–889. https://doi.org/10.1007/s00359-012-0758-6

Orito, K., Chida, Y., Fujisawa, C., Arkwright, P. D., & Matsuda, H. (2004). A new analytical system for quantification scratching behaviourin mice. Br J Dermatol, 150(1), 33–38. https://doi.org/10.1111/j.1365-2133.2004.05744.x

Pagani, M., Albisetti, G. W., Sivakumar, N., Wildner, H., Santello, M., Johannssen, H. C., & Zeilhofer, H. U. (2019). How Gastrin-Releasing Peptide Opens the Spinal Gate for Itch. Neuron, 103(1), 102–117 e105. https://doi.org/10.1016/j.neuron.2019.04.022

Ramirez, F. D., Chen, S., Langan, S. M., Prather, A. A., McCulloch, C. E., Kidd, S. A., Cabana, M. D., Chren, M. M., & Abuabara, K. (2019). Association of Atopic Dermatitis With Sleep Quality in Children. JAMA Pediatr, 173(5), e190025. https://doi.org/10.1001/jamapediatrics.2019.0025

Sachs, B. D. (1988). The development of grooming and its expression in adult animals. Annals of the NY Acad Sci, 525, 1–17. https://doi.org/10.1111/j.1749-6632.1988.tb38591.x

Stumpf, A., Burgmer, M., Schneider, G., Heuft, G., Schmelz, M., Phan, N. Q., Stander, S., & Pfleiderer, B. (2013). Sex differences in itch perception and modulation by distraction--an FMRI pilot study in healthy volunteers. PLoS One, 8(11), e79123. https://doi.org/10.1371/journal.pone.0079123

Takanami, K., Uta, D., Matsuda, K. I., Kawata, M., Carstens, E., Sakamoto, T., & Sakamoto, H. (2021). Estrogens influence female itch sensitivity via the spinal gastrin-releasing peptide receptor neurons. Proc Natl Acad Sci U S A, 118(31). https://doi.org/10.1073/pnas.2103536118

Tominaga, M., Ozawa, S., Tengara, S., Ogawa, H., & Takamori, K. (2007). Intraepidermal nerve fibers increase in dry skin of acetone-treated mice. J Dermatol Sci, 48(2), 103–111. https://doi.org/10.1016/j.jdermsci.2007.06.003

Toyama, S., Moniaga, C. S., Nakae, S., Kurosawa, M., Ogawa, H., Tominaga, M., & Takamori, K. (2021). Regulatory T Cells Exhibit Interleukin-33-Dependent Migratory Behavior during Skin Barrier Disruption. Int J Mol Sci, 22(14). https://doi.org/10.3390/ijms22147443

Wang, X. J. (2002). Probabilistic decision making by slow reverberation in cortical circuits. Neuron, 36(5), 955–968. https://doi.org/10.1016/s0896-6273(02)01092-9

Wimalasena, N. K., Milner, G., Silva, R., Vuong, C., Zhang, Z., Bautista, D. M., & Woolf, C. J. (2021). Dissecting the precise nature of itch-evoked scratching. Neuron, 109(19), 3075–3087 e3072. https://doi.org/10.1016/j.neuron.2021.07.020

Yosipovitch, G., & Bernhard, J. D. (2013). Clinical practice. Chronic pruritus. N Engl J Med, 368(17), 1625–1634. https://doi.org/10.1056/NEJMcp1208814

